# Preparation of physiologically active inside-out vesicles from plant inner mitochondrial membranes

**DOI:** 10.1101/2023.04.27.538533

**Authors:** Leander Ehmke, Gerd Hause, Ralf Bernd Klösgen, Bationa Bennewitz

## Abstract

For many metabolites, the major barrier between cytosol and mitochondrial matrix is the inner envelope membrane of mitochondria, the site of the respiratory electron transport chain. In consequence, it houses numerous transporters which facilitate the controlled exchange of metabolites, ions, and even proteins between these cellular compartments. While their import into the organelle can be studied with isolated mitochondria or mitoplasts, the analysis of their export from the matrix into the intermembrane space or even the cytosol demands for more sophisticated approaches. Among those, inside-out inner membrane vesicles are particularly useful, since they allow the direct presentation of the potential export substrates to the membrane without prior import into the organelle. Here we present a protocol for the isolation of such inside-out vesicles of the inner envelope membrane of plant mitochondria based on repeated freeze/thaw-cycles of freshly prepared mitoplasts. Electron microscopy and Western analysis could show that the majority of the vesicles have single envelope membranes in an inside-out topology. The vesicles are furthermore physiologically active, as demonstrated by assays measuring the enzymatic activities of Complex I (NADH dehydrogenase), Complex V (ATP synthase) and the mitochondrial processing peptidase (MPP) associated with Complex III. Hence, the method presented here provides a good basis for further studies of the inner mitochondrial envelope membrane and mitochondrial export processes.

## Introduction

Plant mitochondria provide a multitude of metabolic reactions and functions which are essential for the cell (e.g., citric acid cycle, respiratory electron transport chain, calcium homeostasis, Fe/S and heme biogenesis, stress adaptation, etc.). The analysis of such processes was strongly supported by the establishment of methods enabling the isolation of intact plant mitochondria and mitochondrial fractions (Day et al., 1985, Gardeström et al., 1978, Moreau and Lance, 1972, Douce et al., 1973, Day et al., 1979). The inner mitochondrial membrane (IMM) is particularly interesting because membrane-bound proteins and complexes are indispensable in many of these reactions and processes, whether as channels, transporters, or enzymes. By osmolysis of intact mitochondria it is possible to disrupt the outer mitochondrial membrane, giving rise to the so-called mitoplasts (Douce et al., 1973), which have the IMM exposed to the surrounding buffer. These mitoplasts thus allow studying many IMM-based processes and properties, though solely from the “outside” of the organelle. In order to gain access to the IMM and its components also from the matrix side, it is necessary to have the membrane inverted. To achieve this, the mitoplasts need to be opened at several sites giving rise to membrane fragments that tend to form sealed vesicles due to their amphiphilic properties. The orientation of these vesicles (inside-out or right-side-out) can be influenced by the method used and the salt concentration of the surrounding medium. For example, French press and low salt medium predominantly lead to right-side-out vesicles, while inside-out vesicles are mostly generated by sonication in high salt buffers (Kay et al., 1985, Møller et al., 1987). Interestingly, the different regions of the IMM, i.e., cristae and inner boundary membrane, each tend to form specific types of vesicles. From cristae regions, relatively large inside-out vesicles are usually obtained, whereas much smaller, right-side-out vesicles are formed from the inner boundary membrane (Møller et al., 1987, Kay et al., 1985).

The first plant IMM vesicles described were generated by sonication of mitochondria isolated from mung bean (*Phaseolus aureus*) (Beyer et al., 1968). Since then, numerous further plant species and tissues have successfully been used (e.g., Rasmusson and Møller, 1991, Braidot et al., 2004). With such vesicles, IMM-based functions like respiratory electron transport, protein topology, enzyme activities, and inhibitors have been studied (Liu et al., 2002, Onda et al., 2007, Korth et al., 1991). However, we have experienced a massive decrease in all examined enzyme activities when using sonication to isolate IMM vesicles, making them unsuitable for our purposes. To better preserve the enzyme functions, the preparation method chosen must thus be as gentle as possible to the proteins and complexes. Therefore, we have adapted a method based on repeated freeze/thaw-cycles using liquid nitrogen that was originally described for the preparation of inside-out vesicles of the plasma membrane (Palmgren et al., 1990). With this method, it was possible to obtain intact and physiologically active inside-out IMM vesicles from pea leave tissue (*Pisum sativum*). We have characterized and validated the vesicles obtained with respect to their morphology, membrane topology and integrity, as well as by determining the enzymatic activities of IMM protein complexes of the respiratory electron transport chain. We think that these inside-out vesicles might be helpful for numerous potential applications, including the analysis of mitochondrial calcium homeostasis or the export of proteins from the matrix to the intermembrane space (IMS).

## Isolation of plant mitochondria and preparation of submitochondrial fractions

### Isolation of mitochondria from pea seedlings

#### Principles

The isolation of intact mitochondria from plant tissue is based on a combination of differential centrifugation and density gradient centrifugation. For more information about the principles of isolating plant mitochondria, see Møller et al. (2021).

#### Solutions

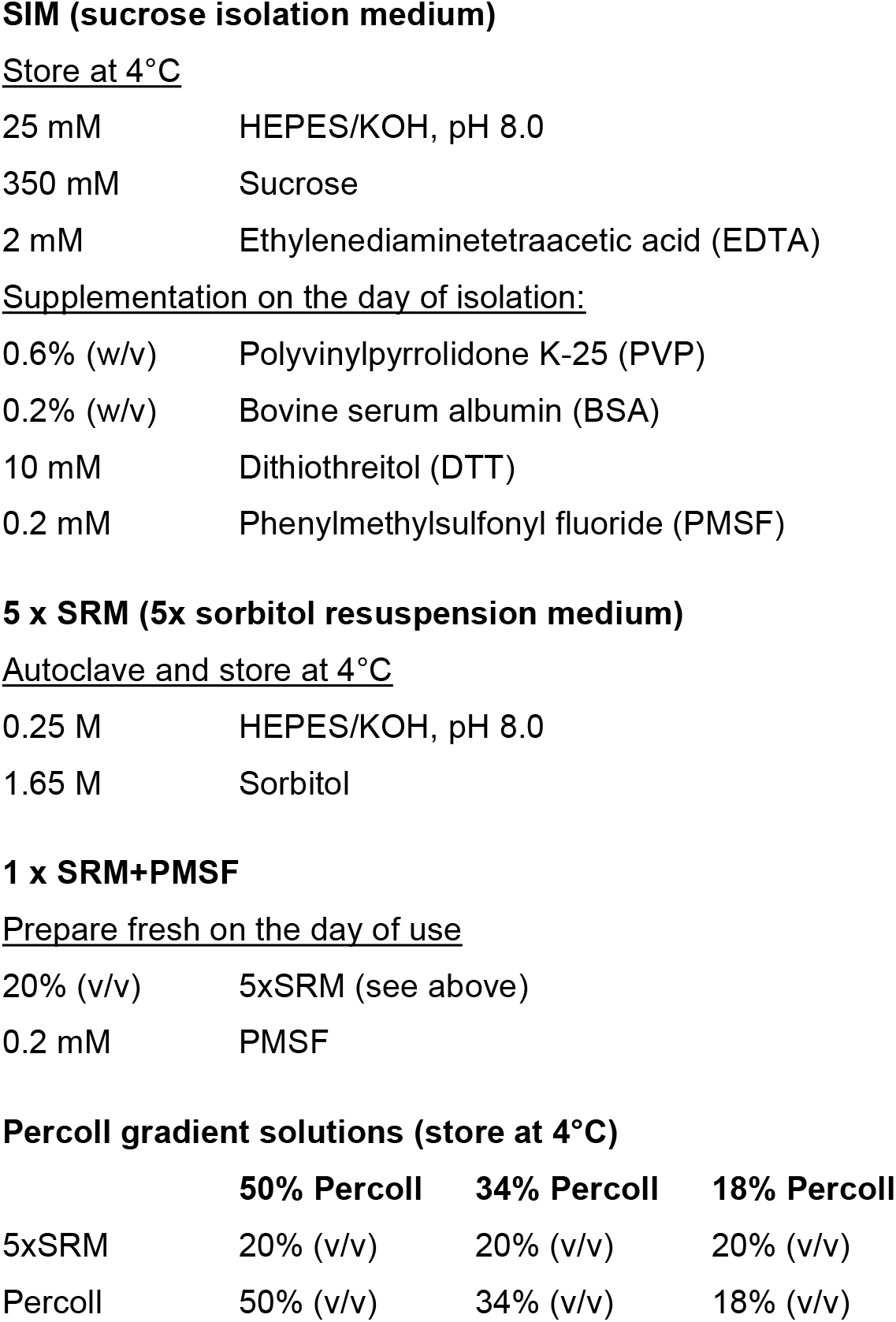

|  |  |
| --- | --- |
| 25 mM | HEPES/KOH, pH 8.0 |
| 350 mM | Sucrose |
| 2 mM | Ethylenediaminetetraacetic acid (EDTA) |
| Supplementation on the day of isolation: |  |
| 0.6% (w/v) | Polyvinylpyrrolidone K-25 (PVP) |
| 0.2% (w/v) | Bovine serum albumin (BSA) |
| 10 mM | Dithiothreitol (DTT) |
| 0.2 mM | Phenylmethylsulfonyl fluoride (PMSF) |
| 0.25 M | HEPES/KOH, pH 8.0 |
| 1.65 M | Sorbitol |
| 20% (v/v) | 5xSRM (see above) |
| 0.2 mM | PMSF |

Percoll gradient solutions (store at 4°C)
|  | 50% Percoll | 34% Percoll | 18% Percoll |
| --- | --- | --- | --- |
| 5xSRM | 20% (v/v) | 20% (v/v) | 20% (v/v) |
| Percoll | 50% (v/v) | 34% (v/v) | 18% (v/v) |

#### Preparatory work

- Cut the nylon mesh (100 µm pore size) and Miracloth (22-25 µm) to size.
- Precool all solutions and equipment to 4°C or on ice.
- Sow pea (e.g., *Pisum sativum* var. Feltham First) and let the seedlings grow for 7–10 d under constant temperature (18–22°C) and light regime (16/8 h light/dark cycle).

##### On the day of isolation

- Finalize the isolation media (SIM, SRM) by adding the supplements.
- Prepare four Percoll step gradients (5 ml 50% Percoll, 8 ml 34% Percoll, 5 ml 18% Percoll) in 30 ml Corex tubes and keep them on ice. ***NOTE***. *To prevent unwanted turbulence, place a flat piece of cork in the Corex tubes onto which the Percoll solutions, and later also the organelle suspension, are pipetted in sequence*.
- Soak soft brush with 1 x SRM+PMSF
- Harvest 80-120 g pea seedlings without roots and keep them on ice until use.

#### Protocol (*timing: 3-4 h*)

The following protocol was modified based on the isolation procedure described by Rödiger et al. (2010). All steps must be performed with precooled solutions and in the cold room or on ice.

1. Homogenize the harvested plant material with approximately 400 ml SIM for 5 × 2 sec in a Waring blender.
2. Filter the homogenate through one layer of nylon mesh and two layers of Miracloth into two 250 ml centrifuge beakers placed on ice. ***NOTE***. *For higher yield, it is advisable to carefully squeeze the filters with the plant debris*.
3. Centrifuge for 5 min at 2,000 *g* (4°C).
4. Carefully pour the supernatant (containing mitochondria) into two new precooled 250 ml centrifuge beakers. ***NOTE***. *The sediment will easily detach from the walls*.
5. Centrifuge for 10 min at 6,000 *g* (4°C).
6. Pour the supernatant (containing mitochondria) into two new precooled 250 ml centrifuge beakers. ***NOTE***. *In order to prevent spillover of the chloroplast-containing sediment, pour the supernatant off in one go but stop as soon as detached sediment fractions approach the rim of the beaker*.
7. Centrifuge for 10 min at 16,000 *g* (4°C).
8. Discard the supernatant while retaining the sediment (containing mitochondria).
9. Use the soft brush to carefully resuspend each sediment in 2 ml 1 x SRM+PMSF. ***NOTE***. *To avoid the formation of lumps during resuspension, the sediment should initially be resuspended in a smaller volume*.
10. Distribute the organelle suspension equally onto the four Percoll gradients.
11. Centrifuge for 45 min at 12,000 *g* (4°C) with low acceleration and deceleration. ***NOTE***. *Intact mitochondria will accumulate in the interphase between 50% and 34% Percoll*.
12. Remove the upper phases with a Pasteur pipette taking care not to take any mitochondria.
13. Carefully collect the interphase containing the mitochondria and transfer it to new precooled 30 ml Corex tubes.
14. Fill the Corex tubes to approximately 25 ml with 1 x SRM+PMSF, mix carefully and centrifuge for 10 min at 12,000 *g* (4°C).
15. Remove and discard approximately 50% of the supernatant.
16. Repeat steps 14 and 15 until a solid sediment has formed and the supernatant remains clear.
17. Discard the supernatant entirely and resuspend the sediment in a few milliliters of 1 x SRM+PMSF.
18. Combine the mitochondrial suspensions in a single Corex tube.
19. Centrifuge for 5 min at 12,000 *g* (4°C).
20. Discard the supernatant, resuspend the sediment in 1 ml 1 x SRM+PMSF and transfer the suspension to a 1.5 ml Eppendorf tube.
21. Centrifuge for 5 min at 16,000 *g* (4°C).
22. Resuspend the sediment in 100 µl 1 x SRM+PMSF.
23. Determine the protein concentration of the mitochondria suspension by Bradford assay. ***ALTERNATIVE***. *Alternatively, a simpler Percoll gradient consisting of 13 ml 34% Percoll and 5 ml 18% Percoll can be used. In this case, the mitochondria are found together with starch granules and other high-density particles at the bottom of the Corex tube after centrifugation. This procedure is faster and usually leads to higher yields at the expense of some impurities with high-density particles*.

### Preparation of mitoplasts and inside-out IMM vesicles

#### Principles

Mitoplasts are prepared from isolated mitochondria by osmolysis in a hypotonic buffer. IMM vesicles are subsequently obtained by repeated freeze/thaw-cycles of mitoplasts resuspended in a buffer containing 20 mM MgCl_2_, which masks membrane charges, allows the cristae regions to stack and thus supports the formation of inside-out vesicles (Kay et al., 1985, Møller et al., 1987) **(Figure 1)**.

**Figure 1.**
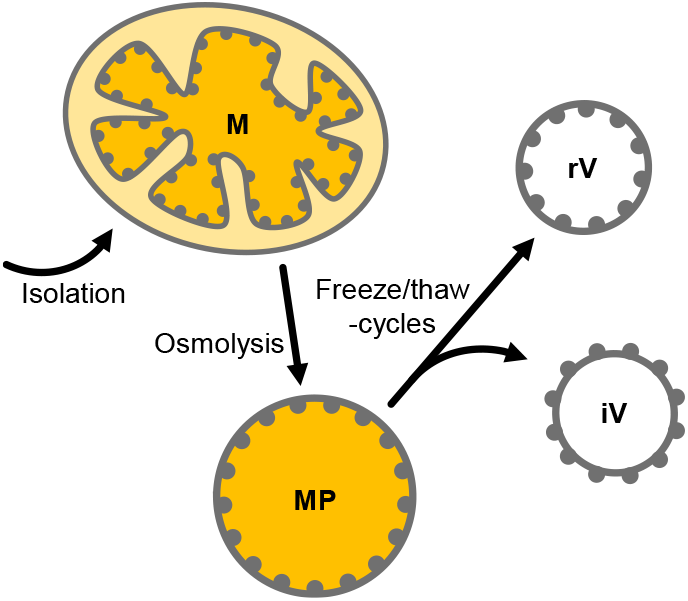
Schematic representation of mitoplast (MP) preparation by osmolysis of mitochondria (M) followed by fractionation into inside-out (iV) and right-side-out (rV) IMM vesicles by repeated freeze/thaw-cycles. Gray dots representing protein complexes attached from the matrix-side to the inner envelope membrane are shown to illustrate the respective membrane orientation.

#### Solutions

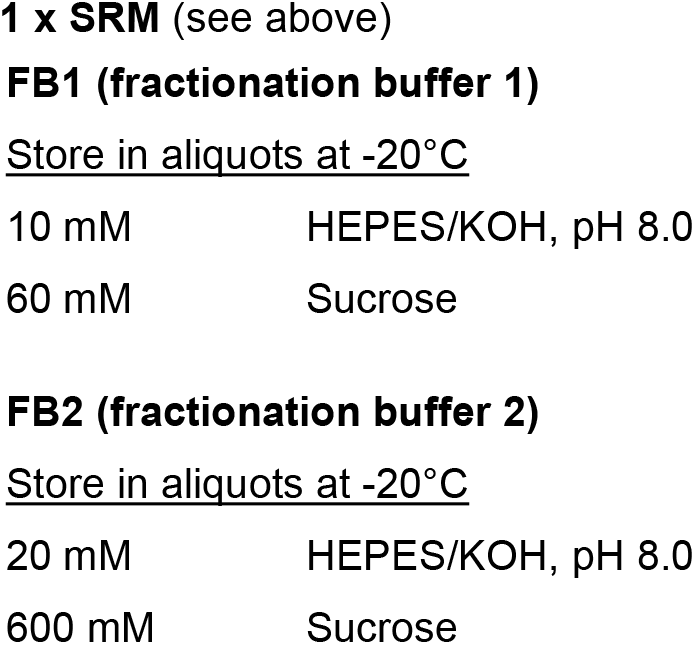

|  |  |
| --- | --- |
| 10 mM | HEPES/KOH, pH 8.0 |
| 60 mM | Sucrose |
| 20 mM | HEPES/KOH, pH 8.0 |
| 600 mM | Sucrose |

#### On the day of isolation

- Supplement FB1 and FB2 with 1mM PMSF each.

#### Protocol (*timing: ∼45 min*)

1. Centrifuge the suspension of isolated mitochondria corresponding to 0.5–1 mg protein for 5 min at 16,000 *g* (4°C).
2. Resuspend the sediment in FB1 (final concentration 10 µg protein/µl).
3. Incubate the suspension for 15 min on ice. ***NOTE***. *During osmolysis, the outer envelope membrane of the mitochondria is disrupted, ultimately leading to mitoplast formation*.
4. Add FB2 in a 1:1 ratio and incubate for 10 min on ice.
5. Centrifuge for 15 min at 16,000 *g* (4°C).
6. Resuspend the sedimented mitoplasts in 1 x SRM (final concentration 10 µg protein/µl).
7. For further fractionation, supplement the assays with MgCl_2_ to 20 mM.
8. Shock-freeze the suspension in liquid nitrogen for 20 sec and let it thaw while holding the tubes in your hands.
9. Repeat step 8 twice. ***NOTE***. *The repeated freezing and thawing of mitoplasts causes the inner envelope membrane to rupture and to reassemble spontaneously. This results in the formation of vesicles, while the presence of MgCl*_*2*_ *supports an inverted membrane orientation (Kay et al*., *1985, Møller et al*., *1987)*.

## Characterization of the IMM vesicles

### Morphology (Electron microscopy)

#### Principles

Transmission electron microscopy (TEM) is a suitable method to determine the morphology, quality, and purity of the isolated vesicles. For this purpose, it is necessary to fix the samples with glutaraldehyde (preservation of ultrastructure) and osmium tetroxide (preservation of ultrastructure and staining of membranes). Then the vesicles are immobilized with Agar, dehydrated with ethanol and infiltrated with an epoxy resin. After polymerization of the resin, the samples are ultra-thin sectioned, transferred to TEM grids, post-stained with heavy metals, and analyzed by electron microscopy.

#### Solutions

|  |  |
| --- | --- |
| 0.2 M | SCB, pH 7.2 |
| 6% (w/v) | Glutaraldehyde (**TOXIC!**) |
| 0.33 M | Sorbitol |

#### Protocol

1. Supplement vesicles corresponding to 100 µg protein with one volume of freshly prepared fixative **(TOXIC!)**, incubate at room temperature for 30–60 min with gentle shaking and store overnight at 4°C for fixation.
2. Centrifuge for 15 min at 22,000 *g* (4°C).
3. Discard the supernatant and resuspend the sediment in 4% Agar (dissolved freshly in 0.1 M SCB) to immobilize the vesicles.
4. Cut the solidified Agar into small pieces and transfer them into a 5 ml sample tube containing 0.1 M SCB.
5. Wash 6 times for 8 min each with 0.1 M SCB.
6. Discard the supernatant and add 1% OsO_4_ (dissolved in 0.1 M SCB).
7. Incubate for 60 min at room temperature.
8. Wash twice for 10 min each with H_2_O.
9. Discard the supernatant and dehydrate the samples stepwise with (a) 10% EtOH, (b) 30% EtOH, and (c) 50% EtOH for 30 min each.
10. Discard the supernatant and add 1% uranyl acetate (dissolved in 70% EtOH).
11. Incubate for 30 min at room temperature. ***NOTE***. *If necessary, the samples can be stored for several days in 70% EtOH at 4°C*.
12. Discard the supernatant and dehydrate the samples further, once with 90% EtOH and twice with 100% EtOH (for 30 min each).
13. Remove the supernatant and incubate the samples stepwise at room temperature as follows:
  a. 25% SER (diluted with 100% EtOH) for 3 h
  b. 50% SER (diluted with 100% EtOH) for 4 h
  c. 75% SER (diluted with 100% EtOH) overnight
14. Discard the supernatant and incubate the samples in 1 ml SER for 6 h at room temperature.
15. Discard the supernatant and incubate the samples in 1 ml SER for 18 h at 70°C to achieve polymerization.
16. Prepare sections from the samples using an ultramicrotome and transfer them to copper grids covered with a Cedukol film (Merck, Darmstadt, Germany).
17. Stain the sections at 25°C for 60 min with 1% uranyl acetate and 10 min with 3% lead citrate using an automatic section stainer.
18. Analyze the sections by TEM. ***NOTE***. *Considering numerous extended incubation steps, the actual TEM analysis usually takes place 8–10 days after starting the procedure*.

#### Results

The electron micrographs of the vesicle fraction reveal the presence of a variety of structures **(Figure 2)**. While most vesicles have single envelope membranes and diameters ranging from 50 nm to 500 nm, also a few vesicles with double envelope membranes as well as membranous fragments are visible. Sporadically, even seemingly intact mitochondria can be found, which are characterized by a double envelope membrane, a darker appearance indicating an electron-dense matrix, and circular membrane structures within the matrix indicative of cross-sections of cristae.

**Figure 2.**
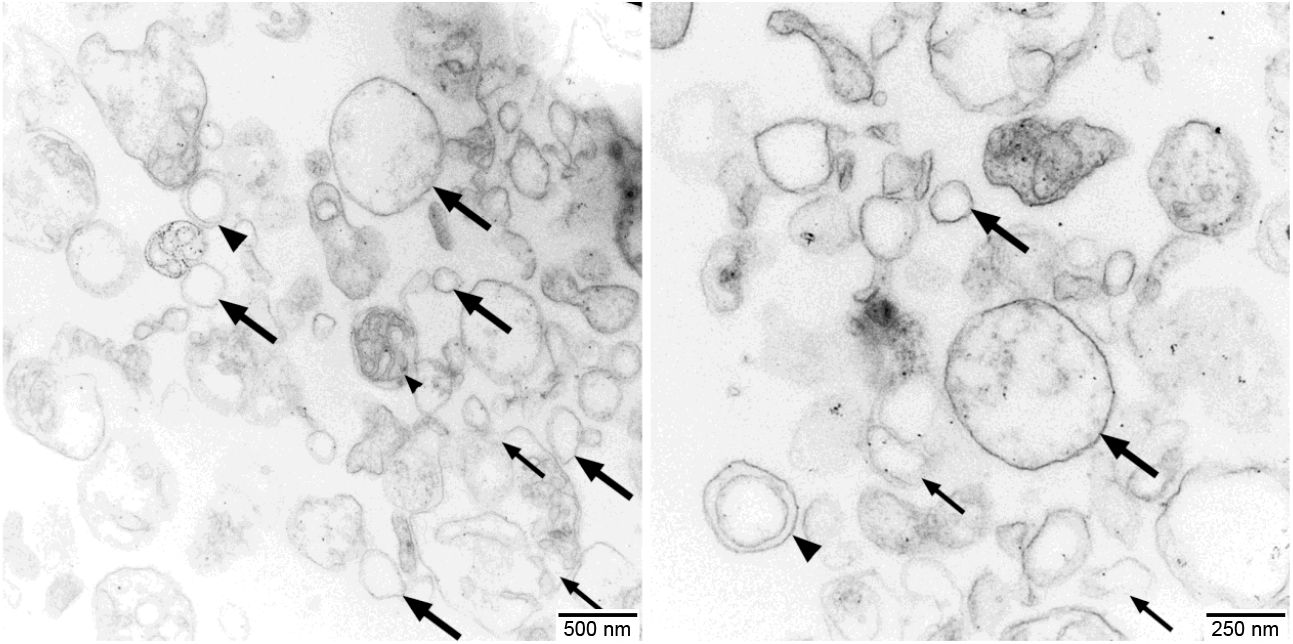
Electron micrographs of the vesicle fractions. Ultrathin sections (70 nm) made with an Ultracut R ultramicrotome (Leica, Wetzlar, Germany) were post-stained with uranyl acetate and lead citrate in an EM-Stain apparatus (Leica) and subsequently analyzed by TEM (Zeiss EM 900, Oberkochen, Germany) operating at 80 kV. Micrographs were taken with an SSCCD SM-1k-120 camera (TRS, Moorenweis, Germany). Single-membrane vesicles (large arrows), double-membrane vesicles (large arrowheads), undefined membrane fragments (small arrows), and a few seemingly intact mitochondria (small arrowheads) are indicated.

### Membrane orientation (Western analysis)

#### Principles

One approach to determine the orientation of the membrane vesicles rests on treatment of such vesicles with highly concentrated solutions of mild chaotropic salts (e.g., 2 M NaBr) to release membrane-attached proteins and protein complexes **(Figure 3A)**. The supernatant and membrane fractions obtained are subsequently analyzed with Western assays employing antibodies against IMM components with known membrane topology.

**Figure 3.**
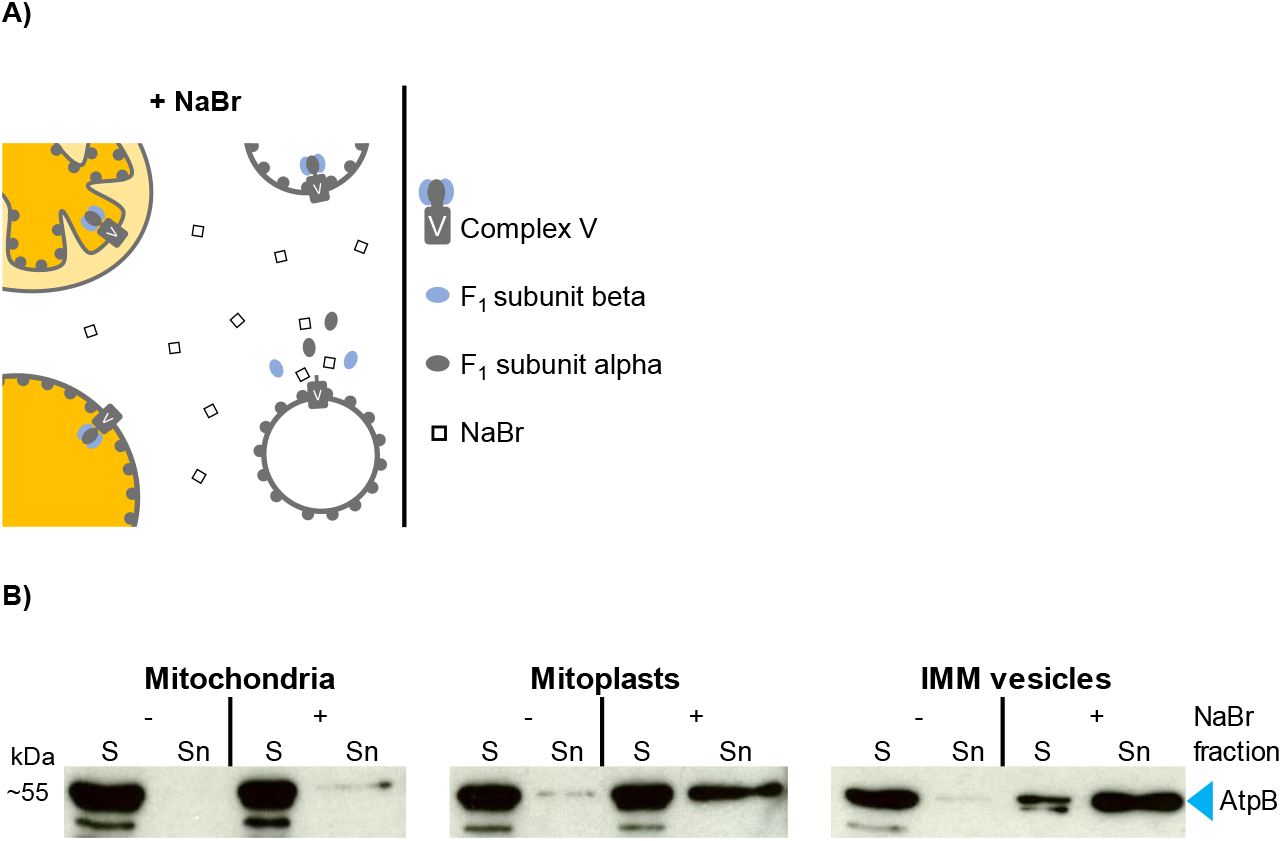
**(A)** Schematic representation of the effect of 2 M NaBr treatment on mitochondria, mitoplasts and IMM vesicles. If accessible to the chaotropic salt, ATP synthase F_1_ subcomplexes are released from the membrane-embedded Fo subcomplex resulting in the accumulation of F_1_ subunits alpha and beta in the supernatant. **(B)** Immunodetection of F_1_ subunit beta (AtpB) in mitochondria, mitoplasts and IMM vesicles. Sediment (S) and supernatant (Sn) fractions obtained after NaBr (+) or mock treatment (–) were examined by Western analysis using antibodies raised against AtpB.

#### Solutions

|  |  |
| --- | --- |
| 250 mM | TRIS/HCl, pH 6.8 |
| 8% (w/v) | SDS |
| 40% (w/v) | Glycerol |
| 20% (w/v) | 2-Mercaptoethanol |
| 0.016% (w/v) | Coomassie G-250 |

#### Protocol (*timing: ∼1 h for steps 1–6*)

1. Collect two aliquots each of mitochondria, mitoplasts and IMM vesicles (each corresponding to 100 µg protein) in Eppendorf tubes and centrifuge at 4°C for either 15 min at 22,000 *g* (vesicles) or 5 min at 16,000 *g* (mitochondria and mitoplasts).
2. Discard the supernatants and carefully resuspend the sediments in either 50 µl 1 x SRM or 50 µl 1 x SRM supplemented with 2 M NaBr (one aliquot per sample each).
3. Incubate on ice for 5 min.
4. Centrifuge at 4°C for either 15 min at 22,000 *g* (vesicles) or 5 min at 16,000 *g* (mitochondria and mitoplasts).
5. Transfer the supernatants to fresh Eppendorf tubes containing 50 µl 4 x LSB each (= Sn fractions, Figure 3B)
6. Resuspend the sediments in 100 µl 2 x LSB each (= S fractions, Figure 3B) ***NOTE***. *Samples can be stored at -20°C until electrophoresis is performed*.
7. Heat the samples for 2 min at 95°C and analyze 5 µl each (corresponding to 5 µg protein starting material) by SDS-PAGE.
8. After electrophoresis, blot the proteins onto a PVDF membrane and perform a standard Western analysis using suitable antibodies.

#### Results

In the experiment shown in **Figure 3B**, we have used antisera against the F_1_ subunit beta of mitochondrial ATP synthase (AtpB), which is exposed to the matrix side of the IMM, to determine the topology of the IMM vesicles prepared. In an inside-out orientation of the vesicles, treatment with 2 M NaBr leads to the dissociation of the F_1_ subcomplex from the membrane-spanning Fo subcomplex of ATP synthase (Complex V) and thus to the accumulation of AtpB in the supernatant **(Figure 3A)** (Hatefi and Hanstein, 1970).

Treatment of isolated intact mitochondria with 2 M NaBr releases only minor amounts of AtpB into the supernatant **(Figure 3B)**, demonstrating that most organelles remain intact during the isolation procedure. In the mitoplast fraction, the proportion of AtpB released is higher, indicating some disruption also of the IMM during osmolysis. Still, the majority of AtpB remains membrane-bound, in line with the assumed robustness of the mitoplasts against osmolysis. In contrast, treatment of the vesicle fraction with 2 M NaBr releases most of the AtpB (>> 50%) into the supernatant, suggesting that the majority of the IMM vesicles have adopted an inside-out orientation during preparation.

### Enzymatic activity of membrane complexes

An alternative approach to evaluate the integrity and orientation of the IMM vesicles rests on determining the enzymatic activity of protein complexes of the respiratory electron transport chain. Here, we have determined the activities of Complex I, Complex V, and the mitochondrial processing peptidase (MPP), which in plant mitochondria is associated with Complex III (Braun et al., 1992, Braun et al., 1995).

### Complex I activity

#### Principles

Complex I activity can be determined by measuring the oxidation of NADH to NAD^+^, which is traced photometrically by the decrease of absorption at its maximum, 340 nm. However, while Complex I provides the predominant NADH oxidizing activity, plant mitochondria have at least three further membrane-bound NADH dehydrogenases. One is found in the outer envelope membrane (Douce et al., 1973) and two in the inner envelope membrane, from which one faces the matrix (Douce et al., 1973, Møller and Palmer, 1982, Rasmusson et al., 2004), whereas the other exposes its active site to the intermembrane space. Thus, all Complex I activity measurements based on NADH oxidation have to take into account the presence of these competing activities.

In the presence of mitoplasts and right-side-out IMM vesicles, solely the IMS-facing NADH dehydrogenase of the inner envelope membrane has access to externally supplied NADH because NADH is neither actively imported into the matrix nor can it diffuse across biomembranes (Jagow and Klingenberg, 1970) **(Figure 4A)**. In the presence of inside-out IMM vesicles, oxidation of NADH is catalyzed by both Complex I and the second NADH dehydrogenase facing the matrix (Møller and Palmer, 1982). Due to the predominant NADH dehydrogenase activity of Complex I, the oxidation rate of externally added NADH should be considerably higher for inside-out IMM vesicles than for any other submitochondrial fraction.

**Figure 4.**
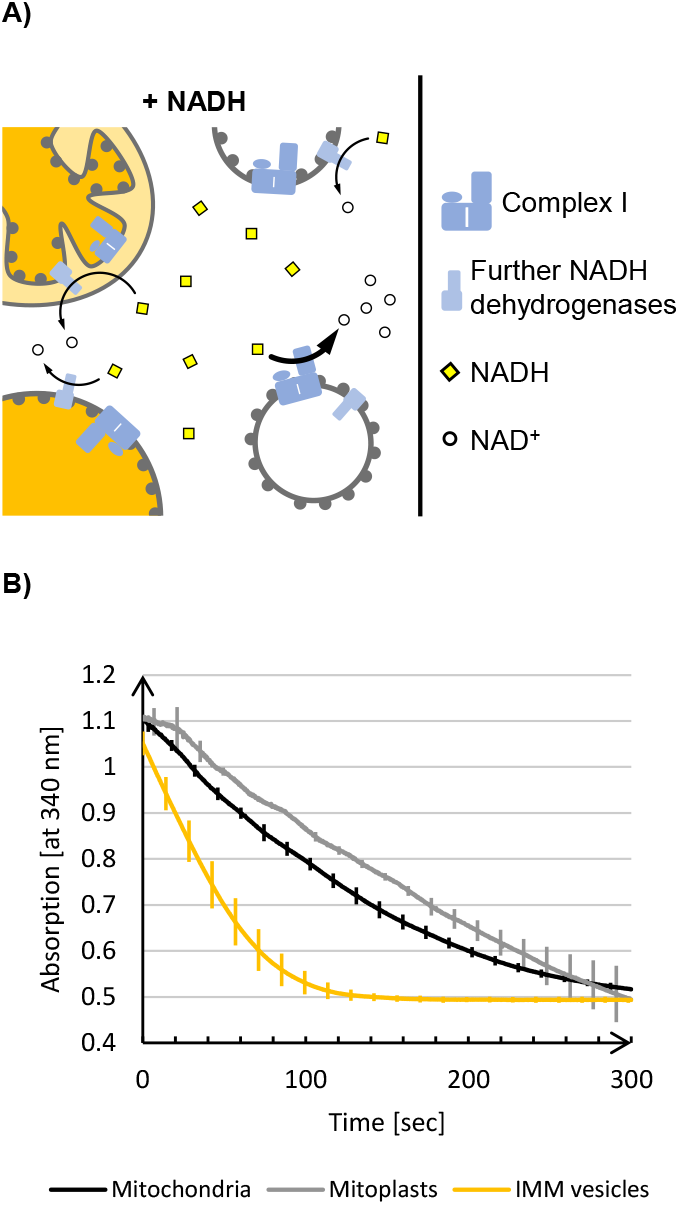
**(A)** Schematic representation of the NADH dehydrogenase activity assay. Intact plant mitochondria, mitoplasts, and right-side-out vesicles can oxidize externally added NADH only via the NADH dehydrogenases facing the intermembrane space, since NADH is not imported into the matrix. In contrast, the inverted membrane orientation of inside-out IMM vesicles allows accessibility of Complex I to the added NADH leading to a massive increase in NADH oxidation. **(B)** Photometric determination of NADH oxidation at 340 nm in mitochondria, mitoplasts and IMM vesicles using a UV-1900 UV-VIS Spectrophotometer (Shimadzu, Kyōto, Japan) and the software UVProbe (v2.70, Shimadzu). Mean values and standard deviations of 3 independent measurements of 5 min each are shown.

In order to exclude any influence of the respiratory electron transport chain on the NADH dehydrogenase activity of Complex I, the assays are supplemented with antimycin A and potassium cyanide (KCN) to inhibit Complex III and Complex IV, respectively. As an electron acceptor, ferricyanide is instead present in excess amounts to prevent any backreaction.

#### Solutions

|  |  |
| --- | --- |
| 20% (v/v) | 5 x SRM (see above) |
| 240 μM | Potassium cyanide (**TOXIC!**) |
| 4 μM | Antimycin A (**TOXIC!**) |

#### Protocol (*timing: ∼7 min per measurement*)

The following protocol was modified based on enzyme activity measurements described by Barrientos et al. (2009) and Gnandt et al. (2017).

1. Add 800 µl 1 x SRM, 200 µl C1 buffer **(TOXIC!)**, and mitochondria, mitoplasts or IMM vesicle suspension corresponding to 150 µg protein into a 1 ml cuvette and measure the blank value at 340 nm.
2. Start the reaction by adding 1.6 µl 100 mM NADH and 10 µl 100 mM ferricyanide.
3. Record the absorption at 340 nm for at least 5 min.

#### Results

When mitochondria or mitoplasts are analyzed, a constant and almost linear decrease in absorption at 340 nm is observed until, after approximately 5 min, the blank value is reached **(Figure 4B)**. In contrast, in the presence of IMM vesicles, such complete oxidation of the NADH added is achieved within approximately 2 min, demonstrating significantly higher NADH dehydrogenase activity due to the exposure of Complex I on the surface of inside-out IMM vesicles.

### Complex V activity

#### Principles

The proton gradient across the inner mitochondrial membrane generated by the respiratory electron transport chain is utilized by mitochondrial Complex V to catalyze the synthesis of ATP. However, the reaction can also proceed backwards at high ATP concentrations, leading to transmembrane proton pumping at the expense of ATP cleavage. In consequence, inside-out IMM vesicles containing active Complex V show internal acidification in the presence of ATP, which can be measured with the help of the basic dye Acridine Orange (AO) **(Figure 5A)**. At neutral pH, AO monomers remain deprotonated and thus membrane permeable, whereas in their protonated form, i.e., at acidic pH, they dimerize and become membrane impermeable. Hence, in the absence of ATP, AO remains largely unaffected, even in the presence of inside-out IMM vesicles. After the addition of ATP, which leads to acidification of the vesicle lumen, AO is protonated, dimerized and thus trapped in the interior of the vesicles.

**Figure 5.**
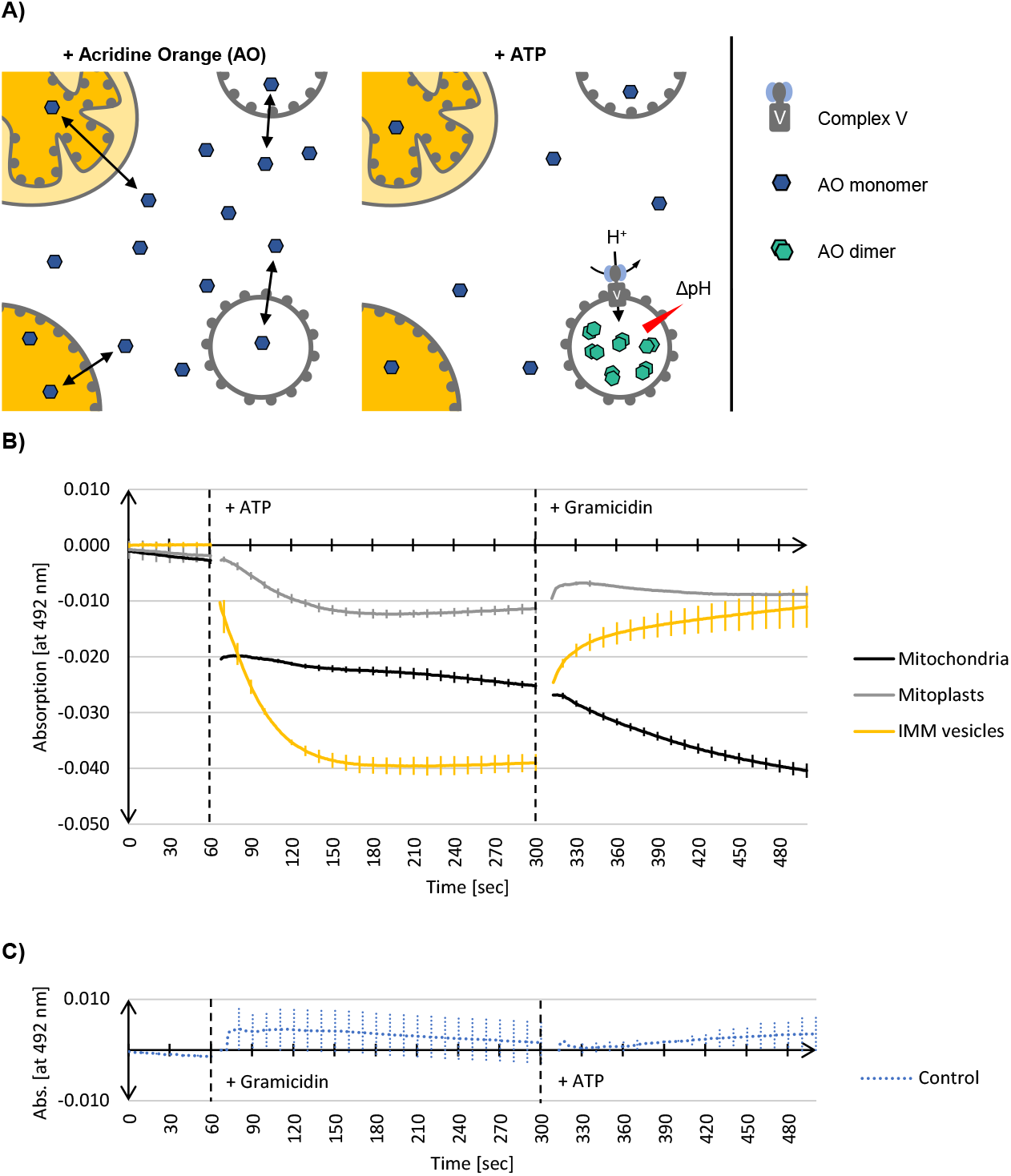
**(A)** Schematic representation of Complex V activity assays (see text for details). **(B)** Photometric determination of the decline of deprotonated Acridine Orange monomer (AO) in the presence of mitochondria, mitoplasts and IMM vesicles. Mean values and standard deviations of 3 independent experiments each are shown. Mitochondria, mitoplasts, or IMM vesicles resuspended in buffer containing AO were placed in a 1 ml cuvette into a UV-1900 UV-VIS Spectrophotometer (Shimadzu, Kyōto, Japan). After 60 sec, the reaction was started by the addition of ATP. After a total of 300 sec, the assays were additionally supplemented with the protonophor Gramicidin. After 500 sec recording was terminated. **(C)** In the control reaction (Control), assays containing IMM vesicles were supplemented after 60 sec with Gramicidin and after a total of 300 sec with ATP. The graphs were generated by the software UVProbe (v2.70, Shimadzu).

The two isoforms of AO can be distinguished photometrically because only monomeric AO shows strong absorption at 492 nm (Falcone et al., 2002). Hence, the decrease of absorption at 492 nm is a measure of the AO dimerization caused by the Complex V-driven acidification of the vesicle lumen. Subsequent supplementation of the assays with Gramicidin, a protonophore disrupting transmembrane proton gradients (Wakiuchi et al., 1988), leads to the deprotonation of AO and the dissociation of the AO dimers, which in consequence results in the recovery of absorption at 492 nm.

This method is suitable not only for determining the orientation of IMM vesicles but provides also information about the integrity of the examined vesicles.

#### Solutions

|  |  |
| --- | --- |
| 25 mM | HEPES/KOH, pH 7.5 |
| 300 mM | Sorbitol |
| 25 mM | KCl |
| 5 mM | MgCl<sub>2</sub> |
| 1 mM | DTT |
| 6 µM | Acridine Orange |

#### Protocol (*timing: ∼15 min per measurement*)

1. Add mitochondria, mitoplasts or vesicles corresponding to 100 µg protein to 1 ml C5 buffer that is preheated to 25°C.
2. Transfer the suspension to a 1 ml cuvette placed in a photometer and incubate for 5 min.
3. Set to zero at 492 nm and record the data for 1 min.
4. Add 50 µl 20 mM ATP, mix thoroughly and continue recording the extinction at 492 nm for 4 min.
5. Add 2 µl 0.5 mM Gramicidin, mix thoroughly and continue recording the extinction at 492 nm for at least 3 min.

#### Results

As shown in **Figure 5B**, adding ATP to inside-out IMM vesicles leads to an immediate decrease in AO absorption at 492 nm. After approximately 2 min, no further decline in absorption can be observed, which indicates that all AO molecules have been protonated and trapped within the vesicles as a consequence of acidification by Complex V activity. Subsequent supplementation of the assays with the protonophor Gramicidin leads to almost complete recovery of AO absorption, demonstrating that protonation of AO was indeed the result of Complex V activity yielding a proton gradient. This was further confirmed in a control reaction in which the vesicles were first supplemented with Gramicidin to avoid the formation of any proton gradient. In this case, the subsequent addition of ATP no longer leads to any change in AO absorption since the vesicle lumen cannot be acidified.

Also in the presence of mitoplasts, a decrease in AO absorption at 492 nm after addition of ATP and its recovery in the presence of Gramicidin is found, though both to a significantly lower extent **(Figure 5B)**. This is in line with the Western data, which shows that during osmolysis of mitochondria some IMMs get disrupted **(Figure 3B)**, resulting in the formation of a few inside-out IMM vesicles. The mitochondria, on the other hand, show a completely different behavior. Adding ATP leads to an immediate drop in AO absorption at 492 nm, which remains almost constant with time, while Gramicidin causes an even further decrease in absorption. Although we cannot provide a final explanation for this phenomenon, it might have to do with the binding properties of AO to DNA molecules, which also results in a change in absorption (Sayed et al., 2016).

### MPP activity

#### Principles

Mitochondrial processing peptidase (MPP) removes presequences from nuclearly encoded mitochondrial precursor proteins after their import into the mitochondrial matrix. In plant mitochondria, MPP is associated with Complex III of the respiratory electron transport chain and exposes its active site to the matrix (Braun et al., 1995). Thus, with inside-out IMM vesicles, MPP activity should be accessible from the surface of the vesicles facilitating the processing of mitochondrial precursor proteins that are present in the surrounding medium **(Figure 6A)**. Both the precursor and the processing products remain accessible though to externally added protease, in contrast to the situation after import into mitochondria.

**Figure 6.**
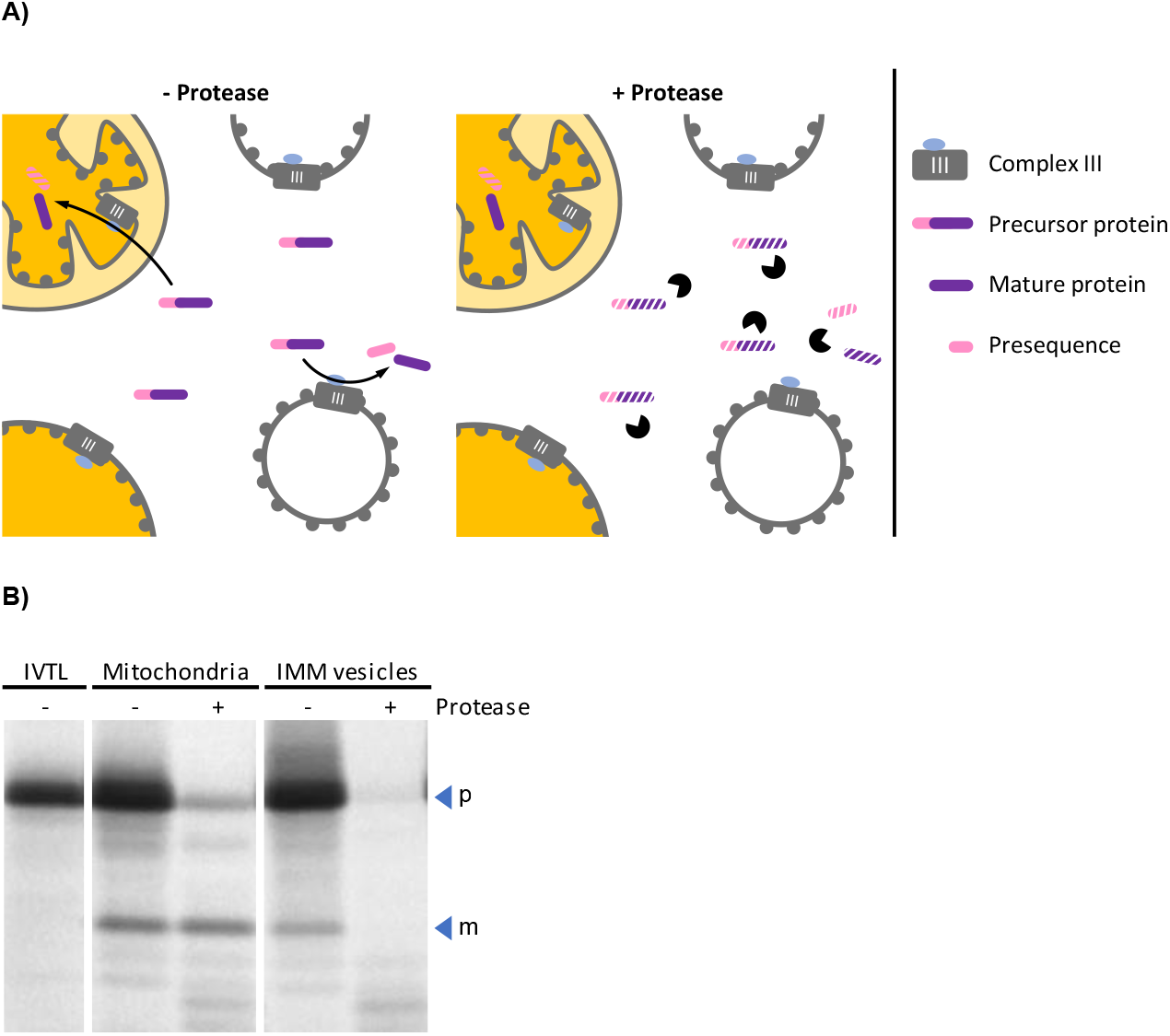
**(A)** Schematic representation of the processes leading to cleavage of mitochondrial precursor proteins by MPP, either after import into mitochondria or by MPP activity exposed on the outer surface of inside-out IMM vesicles (left panel). The two processes can be distinguished from each other by the subsequent addition of protease (right panel), which does not have access to proteins that were imported into the organelle. **(B)** Autoradiogram of an MPP activity assay. The radiolabeled mitochondrial precursor protein obtained by *in vitro* translation in the presence of ^35^S methionine (lane IVTL) was incubated for 20 min at 25°C with either intact mitochondria or IMM vesicles. The samples were reisolated by sedimentation and treated with either protease (+) or mock-treated (–). Stoichiometric amounts of each sample were separated by SDS-PAGE and visualized by phosphorimaging using an FLA-3000 Fluorescence Laser Imaging Scanner (Fujifilm) and the software packages BASReader (v3.14, raytest, Straubenhardt, Germany) and Aida Image Analyzer (v5.0, raytest). The position of precursor (p) and mature (m) protein are indicated by blue arrowheads.

#### Solutions

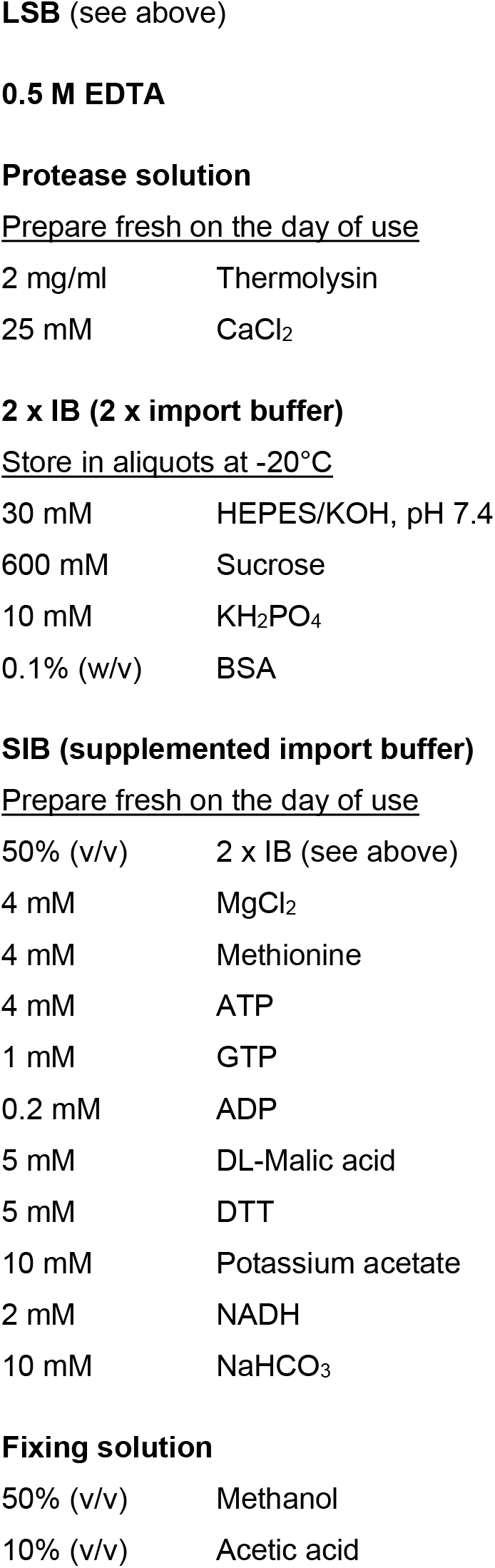

|  |  |
| --- | --- |
| 2 mg/ml | Thermolysin |
| 25 mM | CaCl<sub>2</sub> |
| 30 mM | HEPES/KOH, pH 7.4 |
| 600 mM | Sucrose |
| 10 mM | KH<sub>2</sub>PO<sub>4</sub> |
| 0.1% (w/v) | BSA |
| 50% (v/v) | 2 x IB (see above) |
| 4 mM | MgCl<sub>2</sub> |
| 4 mM | Methionine |
| 4 mM | ATP |
| 1 mM | GTP |
| 0.2 mM | ADP |
| 5 mM | DL-Malic acid |
| 5 mM | DTT |
| 10 mM | Potassium acetate |
| 2 mM | NADH |
| 10 mM | NaHCO<sub>3</sub> |
| **Fixing solution** |  |
| 50% (v/v) | Methanol |
| 10% (v/v) | Acetic acid |

#### Preparatory work

- Prepare radiolabeled mitochondrial precursor proteins by *in vitro* transcription followed by *in vitro* translation in the presence of ^35^S methionine. For details, see Rödiger et al. (2010).

#### Protocol (*timing: ∼1 h 30 min for steps 1–12*)

The following protocol was modified based on the protein transport assays described by Rödiger et al. (2010).

1. Add mitochondria or vesicle suspension corresponding to 100 µg protein and 10 µl *in vitro* translation assay to 90 µl SIB.
2. Incubate for 20 min at 25°C.
3. Centrifuge at 4°C for 5 min at 16,000 *g* (mitochondria) or 15 min at 22,000 *g* (vesicles).
4. Carefully remove supernatant and resuspend each sediment in 200 µl 1 x IB.
5. Divide the samples into two 90 µl aliquots each and discard the rest.
6. While one aliquot of each sample remains untreated, the second aliquot is supplemented with 10 µl protease solution.
7. Incubate for 30 min on ice.
8. Stop protease activity by adding 3 µl 0,5 M EDTA.
9. Centrifuge at 4°C for 5 min at 16,000 *g* (mitochondria) or 15 min at 22,000 *g* (vesicles).
10. Discard supernatant and resuspend each sediment with 100 µl 1 x SRM.
11. Centrifuge at 4°C for 5 min at 16,000 *g* (mitochondria) or 15 min at 22,000 *g* (vesicles).
12. Discard supernatant and resuspend each sediment in 30 µl 2 x LSB.
13. Heat samples for 2 min at 95°C, centrifuge briefly and analyze by SDS-PAGE.
14. After electrophoresis, incubate the gel for 30 min in fixing solution.
15. Dehydrate the gel in vacuum at 80°C maximum (with gradual temperature increase) using a gel dryer (e.g., Bio-Rad Model 583, Hercules, USA). ***NOTE***. *To visualize the position of unlabelled molecular weight marker in the autoradiogram, a small amount of in vitro translation assay in LSB can be dotted with a pipette tip onto the marker bands*.
16. Expose for 0.5–3 days to a phosphor imager screen (e.g., Fujifilm, Minato, Japan). ***NOTE***. *For exposure times exceeding 3 days, the screens should be placed in a radiation-shielding lead chamber to reduce background caused by ambient radiation*. ***NOTE***. *Instead of phosphor imager screens, conventional X-ray films can alternatively be used for exposure. However, due to the lower sensitivity of such films, exposure times will probably be considerably longer*.

## Results

In the experiment shown in **Figure 6B**, the radiolabeled mitochondrial precursor protein is processed to its mature form in the presence of both, intact isolated mitochondria and inside-out IMM vesicles. After import into the mitochondrial matrix, the mature protein is protected by the organellar envelope membranes against externally added protease. In contrast, it remains accessible to the protease if the reaction is instead performed with inside-out IMM vesicles.

## Acknowledgement

This work was supported by a grant from the Deutsche Forschungsgemeinschaft (GRK2498 - 400681449).

